# Spatial segregation and cooperation in radially expanding microbial colonies under antibiotic stress

**DOI:** 10.1101/2020.02.18.954644

**Authors:** Anupama Sharma, Kevin B. Wood

## Abstract

Antibiotic resistance in microbial communities reflects a combination of processes operating at different scales. The molecular mechanisms underlying antibiotic resistance are increasingly understood, but less is known about how these molecular events give rise to spatiotemporal behavior on longer length scales. In this work, we investigate the population dynamics of bacterial colonies comprised of drug-resistant and drug-sensitive cells undergoing range expansion under antibiotic stress. Using the opportunistic pathogen *E. faecalis* with plasmid-encoded (*β*-lactamase) resistance as a model system, we track colony expansion dynamics and visualize spatial pattern formation in fluorescently labeled populations exposed to ampicillin, a commonly-used *β*-lactam antibiotic. We find that the radial expansion rate of mixed communities is approximately constant over a wide range of drug concentrations and initial population compositions. Fluorescence imaging of the final populations shows that resistance to ampicillin is cooperative, with sensitive cells surviving in the presence of resistant cells even at drug concentrations lethal to sensitive-only communities. Furthermore, despite the relative invariance of expansion rate across conditions, the populations exhibit a diverse range of spatial segregation patterns, with both the spatial structure and the population composition depending on drug concentration, initial composition, and initial population size. Agent based models indicate that the observed dynamics are consistent with long-range cooperation, despite the fact that *β*-lactamase remains cell-associated in *E. faecalis*, and experiments confirm that resistant colonies provide a protective effect to sensitive cells on length scales multiple times the size of a single colony. Furthermore, in the limit of small inoculum sizes, we experimentally show that populations seeded with (on average) no more than a single resistant cell can produce mixed communities in the presence of drug. While biophysical models of diffusion-limited drug degradation suggest that individual resistant cells offer only short-range protection to neighboring sensitive cells, we show that long-range protection may arise from synergistic effects of multiple resistant cells, even when they represent only a small fraction of a colony’s surface area. Our results suggest that *β*-lactam resistance can be cooperative even in spatially extended systems where genetic segregation typically disfavors exploitation of locally produced public goods.

## INTRODUCTION

Antibiotic resistance is increasingly viewed as a long-term threat to global health (1, 2). Thanks to decades of seminal breakthroughs in microbial genetics and molecular biology, the molecular-scale conduits of resistance are often well-understood (3), though the diversity and ubiquity of these defense systems paints a dauntingly complex picture of microbial adaptability. In addition, antibiotic resistance can reflect collective phenomena (4, 5, 6, 7, 8, 9, 10, 11, 12, 13, 14, 15, 16, 17, 18, 19, 20), leading to microbial communities that are significantly more resilient than the individual constituent cells. As a result, there is a growing need to understand dynamics of resistance–and of microbial populations in general–across length scales.

In nature bacterial populations frequently exist as spatially structured biofilm populations. Biofilms exhibit a fascinating array of spatiotemporal behavior borne from a combination of heterogeneity, competition, and collective dynamics across length scales (21, 22, 23, 24, 25, 26, 27, 28, 29, 30, 31, 32, 33). Biofilms also contribute to a wide range of clinical complications, including infections of heart valves and surgical implants (34, 35). Work over the last 20 years, in particular, has dramatically improved our understanding of spatial segregation, selection, cooperation, and genetic drift in microbial communities (36, 37, 38, 39, 40, 41, 42, 43, 44, 45, 46, 47, 48, 49, 50, 33, 51, 52, 53, 54, 55). These dynamics often reflect locally heterogeneous cell-cell interactions which lead to emergent behavior on longer length scales. The spatial structure also impacts selection by limiting the competition to a local scale, where a cell only competes with a sub-population of cells in its neighborhood. Further, selection acts differently in expanding biofilms because only the cells at the front contribute to population growth, effectively reducing the population size and enhancing genetic drift. During range expansions of bacterial colonies, the enhanced genetic drift at the growing front frequently gives rise to large spatial sectors of cells from a common lineage (37), though range expansions can either maintain or decrease population diversity depending on the nature and scale of intercellular interactions (37, 43, 44, 42, 56, 57, 49).

Despite an increasingly mature understanding of spatial dynamics in microbial communities, much of our knowledge of antibiotic resistance is derived from experiments in homogeneous (well-mixed) liquid cultures. However, numerous studies indicate that spatial heterogeneity in drug concentration can modulate the evolution of resistance (58, 59, 60, 61, 62, 63, 64, 65). More generally, spatial structure can play an important role in the presence of cooperative resistance mechanisms. When resistance is conferred by drug-targeting enzymes that reduce local antibiotic concentration, for example, the dynamics in even well-mixed communities can be complex and counterintuitive (5, 66, 67, 68, 11, 69, 14, 70). Spatial structure can enhance or inhibit these effective intracellular interactions, creating an environment where cooperation, competition, and evolutionary selection are potentially shaped by spatial constraints (71, 69, 72, 18, 73, 74, 75, 76). For example, microbiologists have long reported the existence of (sometimes annoying (77)) “satellite” colonies (78), drug-sensitive populations that appear near enzyme-producing resistant colonies on agar plates, suggesting that resistance can be cooperative on scales of millimeters or more. Similarly long-range effects have been recently measured for oxygen gradients in biofilms (79), yet metabolic gradients have been shown to be relatively short-ranged (80), and droplet-based printing techniques have shown that specific microscale structure determines dynamics in interacting colicin-producing strains (81). In systems where intercellular interactions are primarily short-range–with resistance driven by intracellular drug degradation, for example (14)–recent work shows that additional physiological mechanisms, such as persistence, might be needed to drive cooperative resistance (18). As a whole, these studies indicate that the length scales that determine multicellular interactions are variable and may depend sensitively on the ecological and biological details of each system. As a result, the length scales dictating cooperative drug resistance, and the biophysical parameters that affect them, are not well understood, leading to a gap in our understanding of how interactions driven by resistance mechanisms shape the dynamics and evolution of bacterial populations.

In this work, we take a step towards closing this knowledge gap by probing the population dynamics of mixed bacterial colonies comprised of drug-resistant and drug-sensitive cells undergoing range expansion under antibiotic stress. As a model system, we use *E. faecalis*, a Gram-positive opportunistic pathogen (82) that readily forms biofilms associated with clinical infections (83, 84, 85). Specifically, we track colony expansion dynamics and visualize spatial pattern formation in fluorescently labeled populations exposed to ampicillin, a commonly-used *β*-lactam antibiotic. Populations are comprised of two strains mixed in varying proportions; one strain is sensitive to ampicillin, while the other is engineered to constitutively express plasmid-encoded *β*-lactamase, an enzyme which confers ampicillin resistance by hydrolyzing the drug’s *β*-lactam ring (86, 87, 88, 89). *β*-lactamase expression in *E. faecalis* is typically constitutive and relatively low-level–at least when compared with expression in *S. aureus*, which harbor an identical enzyme (88). In addition, the enzyme has been isolated from cell pellets, but not from cell-free media, indicating that the enzyme remains cell-associated in *E. faecalis* populations rather than being released into the extracellular media (88). These findings suggest that cooperative resistance may be limited to relatively short length scales, especially in spatially structured populations. Surprisingly, however, we find that the radial expansion rate of mixed communities is approximately constant over a wide range of drug concentrations and (initial) population compositions. Microscopic imaging of the final populations shows that resistance to ampicillin is cooperative, with sensitive cells surviving in the presence of resistant cells even at drug concentrations lethal to sensitive-only communities. Furthermore, despite the relative invariance of expansion rate across conditions, the populations exhibit a diverse range of spatial segregation patterns, with both the spatial structure and the population composition depending on drug concentration, initial composition, and initial population size. Simple mathematical models indicate that the observed dynamics are consistent with global cooperation, and experiments confirm that resistant subpopulations–at times originating from only a single resistant cell–provide a protective effect to sensitive cells on length scales multiple times the size of a single colony.

## MATERIALS AND METHODS

### Bacterial strains and growth conditions

Experiments were performed with *E. faecalis* strain OG1RF. Ampicillin resistant strains were engineered as described in (70). Briefly, we transformed (90) OG1RF with a modified version of the multicopy plasmid pBSU101, which was originally developed as a fluorescent reporter for Gram-positive bacteria (91). The modified plasmid, named pBSU101-BFP-BL, expresses BFP (rather than GFP in the original plasmid) and also constitutively expresses β-lactamase driven by a native promoter isolated from the chromosome of clinical strain CH19 (87). The β-lactamase gene and reporter are similar to those found in other isolates of enterococci and streptococci (86, 92). Similarly, sensitive strains were transformed with a similar plasmid, pBSU101-DasherGFP, a pBSU101 derivative that lacks the β-lactamase insert and where eGFP is replaced by a brighter synthetic GFP (Dasher-GFP; ATUM ProteinPaintbox, https://www.atum.bio/). The reporter plasmids are described in detail in (93, 70). The plasmids also express a streptomycin resistance gene, and all media was therefore supplemented with streptomycin.

### Antibiotics

Antibiotics used in this study included Spectinomycin Sulfate (MP Biomedicals) and Ampicillin Sodium Salt (Fisher).

### Colony growth experiment

Colony-growth experiments were conducted on 1% agar plates with Brain-Heart Infusion media (Remel) at 37°C. Plates were poured 1 day before inoculation and stored in the refrigerator at 4°C. Cells of both strains were grown at 37°C overnight in liquid BHI and spectinomycin. On the day of inoculation on agar plates, the overnight culture was diluted in fresh media and spectinomycin, and was grown for 3 hrs to ensure that the cells are in the exponential phase. 1 mL of this diluted culture was spun down for 3 min at 6800 × *g* and re-suspended in 1 mL of NFW. The densities of such suspensions were measured and adjusted to OD_600_ of 0.1 by adding or removing NFW as needed. Thereafter, the suspensions were mixed in the desired ratios and a droplet of volume 2 *μ L* (approx. 16 × 10^5^ cells) was pipetted onto the agar plate. These plates were then placed in a warm room at 37C to allow the range expansion, which was tracked using a commercial document scanner upto seven days. The experimental reproducibility of results was checked 4 times on different days.

### Image acquisition and analysis

A commercial document scanner controlled by a computer program was used to take time series of images for seven days (94). These images were then processed and used to estimate the colony growth rate and appearance time using custom analysis scripts based in Matlab. At the end of the expansion experiment (marked by no change in colony radius), the colonies were visualized under a fluorescence stereoscope (Olympus SZX16). The red and green fluorescent channel were imaged separately with the exposure time set by the the software accompanying the stereoscope. The acquired images were processed to remove noise and the image overlays were created using Fiji/ImageJ (95). A Python script was used for the quantification of images by creating a binary image for each channel (red or green) using Otsu’s method for thresholding. Thereafter, number of bright pixel were calculated for each channel to estimate the fraction of cells in the final composition. Population compositions measured in this way are not overly sensitive to the specific value of the threshold parameter (see Figure S3) and are expected to well approximate true population composition when subpopulations are well segregated but provide only qualitative estimates of true composition in well-mixed regions.

### Cooperative growth model

To simulate the colony growth dynamics of a mixed microbial population, we developed a simple agent based model of microbial range expansions, similar to the Eden growth model (96). Our goal is not to produce the fine-scale quantitative features of the observed spatial patterns, which may require more detailed models. Instead, our aim was to develop a minimal model to probe the qualitative features of our experiments, most notably the observations that 1) population composition, but not expansion velocity, depends on drug concentrations and initial resistant fraction and 2) spatial patterns are consistent with long-range cooperative resistance.

We begin with an initial population consisting *N*_0_ individual cells arranged within a circle of radius *r* on a two-dimensional lattice. This initial population consists of two subpopulations representing drug-sensitive cells (labeled by −1) and drug-resistant cells (labeled by +1). Each lattice site is occupied by only one type of allele and unoccupied sites are represented by 0. At each time step, a random cell is selected for division, which populates one of the unoccupied neighbouring sites (8 nearest neighbors, in total) by a cell of the same type (∅ → ±1). To model growth in super-MIC drug concentrations, sensitive cells are also allowed to die (−1 → ∅) with probability *d*, which depends on drug concentration. For simplicity, we assume *d* is proportional to AMP concentration, with the proportionality constant chosen so that sensitive-only populations cannot survive at AMP concentrations greater than the MIC.

To incorporate cooperative resistance between resistant and sensitive cells, we assume that the death probability *d* for sensitive cells depends on the local population of resistant cells. In the local version of the model, the death rate depends only on the fraction of nearest neighbor cells that are resistant, while the global version depends on the fraction of resistant cells in the entire population. For simplicity, we assume a simple threshold model where death of sensitive cells occurs at a rate d when the fraction of resistant cells is small but drops to 0 (i.e. the sensitive cell acts like a resistant cell) when the resistant fraction crosses some threshold *f_crit_*. Because this threshold may also depend on drug concentration–intuitively, larger drug concentrations would require larger populations of resistant cells to degrade drug to sub-MIC levels–we also take the threshold to be proportional to AMP concentration.

## RESULTS

To investigate the dynamics of *E. faecalis* populations exposed to β-lactams, we used a previously engineered drug resistant *E. faecalis* strain that contains a multicopy plasmid that constitutively expresses β-lactamase and also expresses a green (GFP-derived) fluorescent marker (Materials and Methods). Sensitive cells harbored a similar plasmid that lacks the β-lactamase insert and expresses a variant of RFP. We grew mixed populations containing sensitive and resistant cells on BHI-agar plates supplemented with various concentrations of antibiotic. We tracked colony growth for 6-7 days post-inoculation using a commercial document scanner, which allowed to us estimate radial expansion velocity and colony appearance time (94), and we then imaged the mixed population at the final time point using fluorescence microscopy (Figure 1(a)) and quantified the relative proportion of sensitive and resistant populations (Materials and Methods).

### Expansion velocity but not population composition is invariant across drug concentrations and initial resistance fractions

Surprisingly, we found that the expansion rate of mixed populations was largely insensitive to initial population composition and drug concentration, including at concentrations that exceed the minimum inhibitory concentration (MIC) for sensitive only populations (Figure 1b; we do note, however, a weak concentration dependence of velocity at low inoculum densities, Figure S1). By contrast, the appearance time of the colonies–that is, the time at which the colony is first detected on the scanner–increases at super-MIC concentrations when the initial resistant fraction is sufficiently small (1 percent). Interestingly, the mixed communities contain sensitive cells following growth in ampicillin that would be lethal to sensitive-only populations (Figure 1c), suggesting that resistance is cooperative, with resistant cells offering a protective effect to sensitive cells. However, the composition of the final population can depend significantly on both drug concentration and initial resistant fraction (Figure 1d).

**FIG 1.**
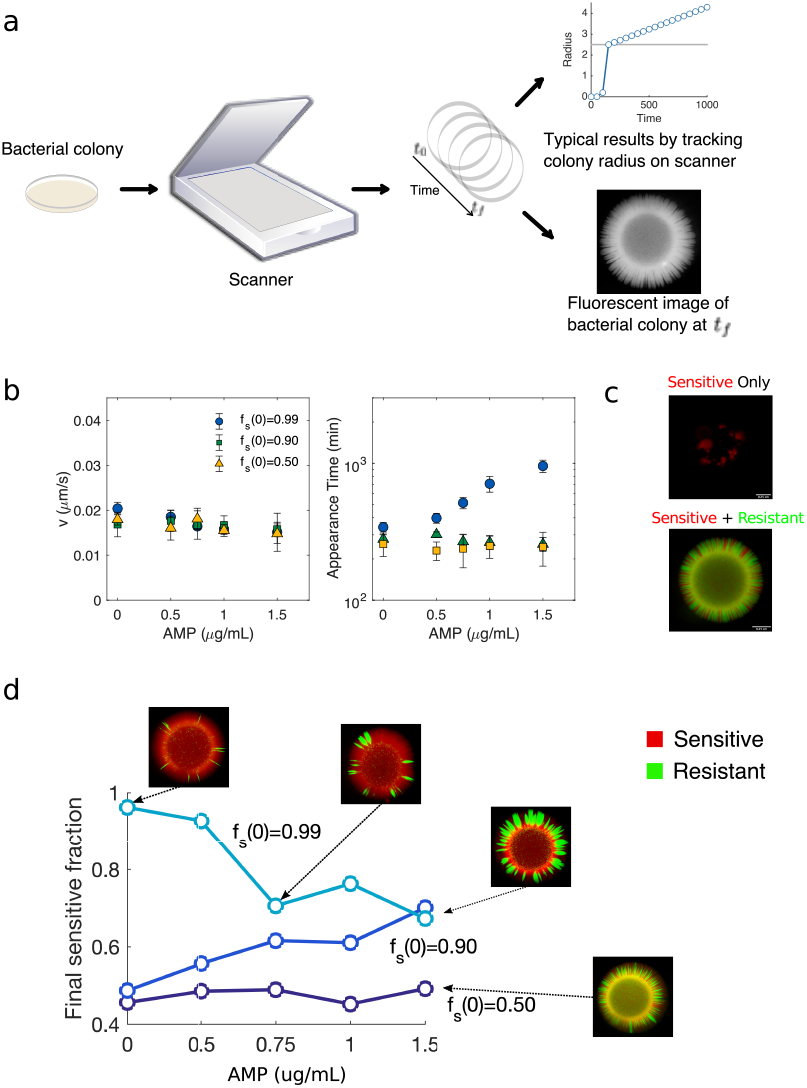
Varying drug concentration and initial resistance fraction alters population composition and appearance time but not expansion velocity. a. Schematic of experimental setup. Colonies are grown on BHI-agar plates and imaged over time using a computer controlled document scanner, allowing for an estimate of the radial expansion velocity and appearance time. At the final time point, the colony is imaged using a fluorescence microscope. b. Expansion velocity (left) and appearance time (right) of mixed colonies containing drug sensitive (WT) and drug resistant (*β*-lactamase producing) strains at different starting ratios (sensitive fraction *f_s_*(0) = 0.99 (circles), 0.90 (squares), or 0.50 (triangles)) at different ampicillin (AMP) concentrations. c. A sample fluorescence microscopy image of a sensitive only (*f_s_*(0) = 1) and a mixed population after growth at super-inhibitory concentrations of AMP (1.5 μg/mL). d. Final sensitive fraction as a function of AMP for populations starting with initial sensitive fractions 0.99 (light blue), 0.90 (blue), 0.50 (dark blue). Images, clockwise from upper right: AMP=0, *f_s_*(0) = 0.99; AMP=0.5, *f_s_*(0) = 0.99; AMP=1.5, *f_s_*(0) = 0.99; AMP=1.5, *f_s_*(0) = 0.50. All colonies were started from 2 *μ*L of population standardized to a density of OD=0.1 (the inoculum contained approximately 2 × 10^5^ cells).

### Observed spatial patterns are consistent with long-range cooperative resistance

To better understand the experimental observations, we developed a simple agent-based model for microbial range expansions in populations of sensitive and resistant cells (Materials and Methods). Briefly, the model assumes cell division-driven expansion along with drug concentration-dependent cell death (in sensitive, but not resistant, cells). In the absence of drug, simulations show the expected genetic segregation characteristic of range expansions (Figure 2a). When exposed to super-inhibitory concentrations of antibiotic, resistant cells preferentially survive, leading to homogeneous populations inconsistent with experimental observations (Figure 2b). To incorporate cooperative resistance, we modified the model so that the rate of cell death of sensitive cells is dependent on either 1) the number of resistant cells in the local neighborhood (“local model”) or 2) the number of resistant cells in the entire community (“global model”). Simulations of the local model reveal co-existence of sensitive and resistant cells in the homeland region where the initial inoculum was placed, but expansion to larger regions is dominated by resistant cells (Figure 2c), a finding that is again inconsistent with observed patterns. By contrast, the global model produces spatial segregation similar to those produced in the absence of drug (Figure 2d). Consistent with experiment, we also find that the globally coupled model leads to expansion velocities invariant across drug concentrations and initial resistant fractions, though the final population composition depends on both factors. Taken together, these simulations suggest that the experimental population dynamics are driven by long-range cooperative resistance between resistant and sensitive cells.

**FIG 2.**
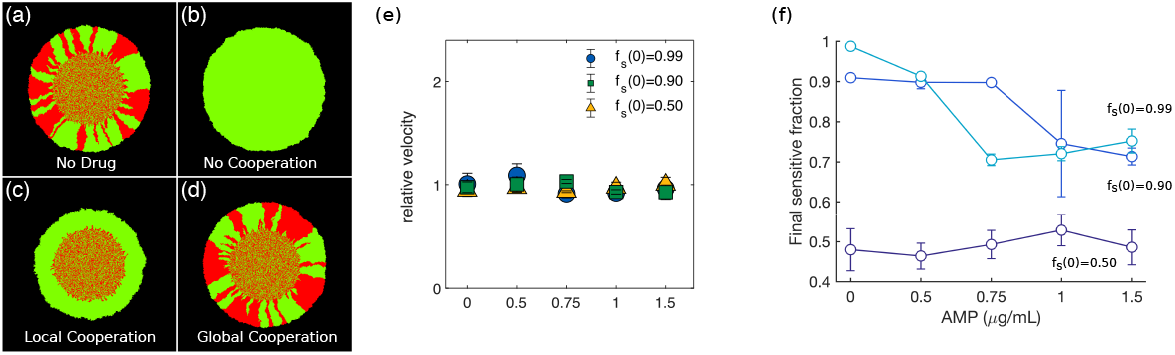
Simulations show long-range cooperative resistance leads to invariant expansion velocity, spatial patterns, and drug-dependent population composition. (a) Simulated colony consisting of drug-sensitive (red) and drug-resistant cells in the absence of drug. (b) In the absence of cooperation, sensitive cells cannot survive at super-inhibitory drug concentrations. (c) Locally cooperative drug resistance, where resistant cells protect neighboring drug sensitive cells from drug-induced cell death, leads to co-existence in the homeland region but drug sensitive cells are not able to survive in the peripheral expanding region. (d) Long-range cooperative resistance, where sensitive cells can survive when the global fraction of resistant cells eclipses a critical threshold, produce spatial patterns similar to those in the absence of drug. Long-range cooperative resistance leads to populations with drug-independent expansion velocity (e) and compositions that depend on drug concentration and initial resistance (f). Expansion velocity is normalized to 1 in the case of no drug.

To investigate the nature of the spatial patterns over a wider range of parameters, we grew and imaged colonies starting from initial resistant fractions as small as 10^−5^ for five different concentrations of ampicillin, including both sub- and super-MIC concentrations. In the absence of drug, colonies starting at the lowest resistant fractions contain no visible resistant cells, though all colonies grown above the MIC contain at least one visible resistant subpopulation. We find that the range expansions typically give rise to the smooth wavefronts and sharp domain boundaries that rarely collide and annihilate (Figure 3), which may be partially due to morphology (46, 47) and non-motile (97) nature of *E. faecalis*. Moreover, in the limit of small initial resistant fraction and super-MIC drug concentrations, two out of the three populations are characterized by a single localized region of resistant cells within a largely sensitive population. These patterns are consistent with long-range cooperative interactions and suggest that the protective effects of resistant cells extend for nearly the entire length of the colony. In this scenario, survival of the population requires a sufficiently large subpopulation of resistant cells–in the extreme case when initial resistant fraction is 10^−5^, initial populations contain fewer than 10 resistant cells, on average–though in the final population these subpopulations need not be spatially distributed (as would be required if cooperation were strictly local). Throughout this regime, final composition is dependent on both initial resistant fraction and drug concentration, and these qualitative trends are also captured by the simulations (Figure 4a-b).

**FIG 3.**
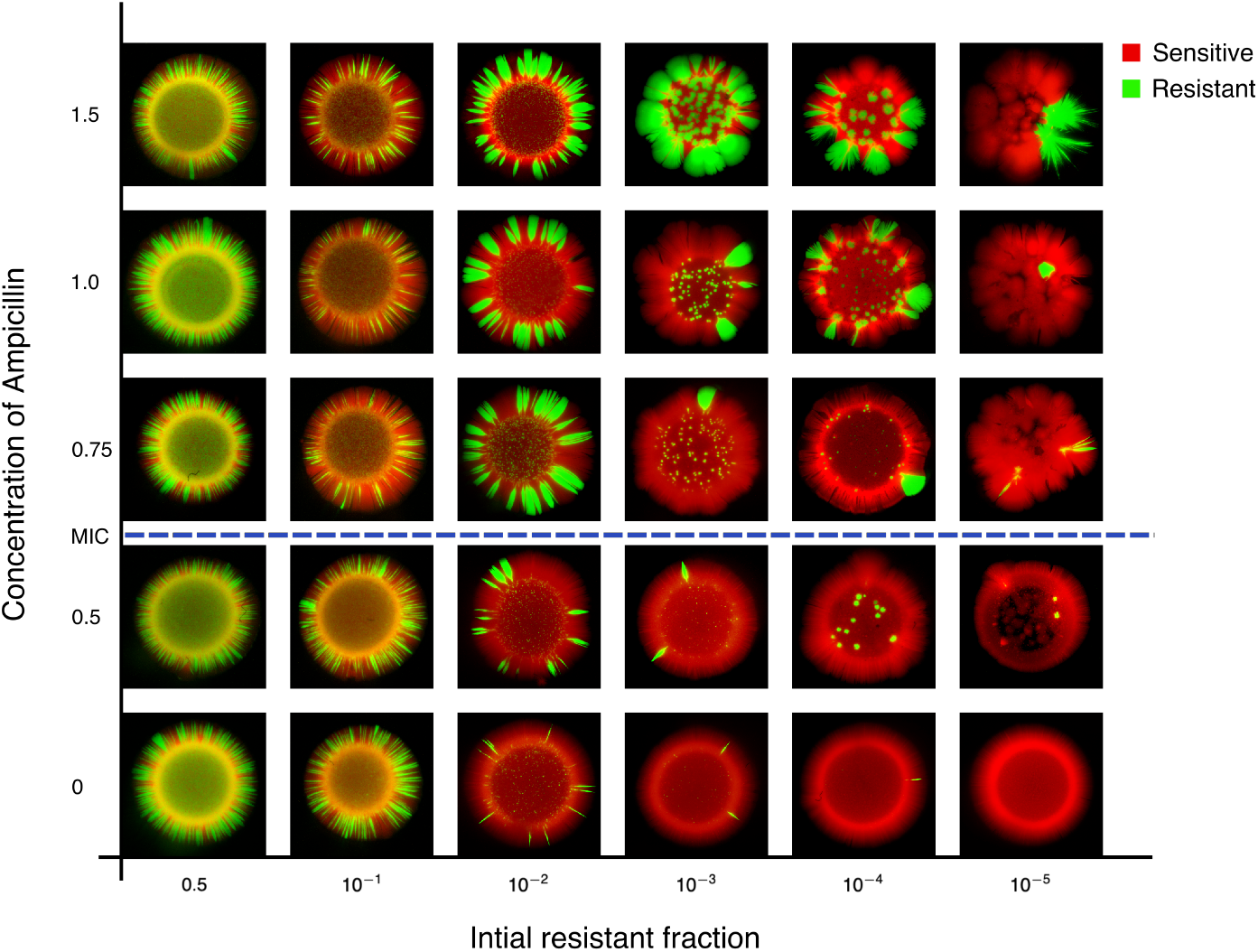
Spatial patterns depend on drug concentration and initial resistant fraction. Images of spatially extended colonies for different concentrations of AMP (vertical axis) and different initial compositions (sensitive fraction increases left to right). Sensitive (red, RFP) and resistant strains (green, GFP) constitutively express different fluorescent proteins. Dashed line indicates the approximate minimum inhibitory concentration (MIC); when drug concentration exceeds the MIC, sensitive-only populations are unable to form colonies (see, for example, Figure 1C). All colonies were started from 2 *μ*L of population standardized to a density of OD=0.1 (the inoculum contained approximately 2 × 10^5^ cells).

**FIG 4.**
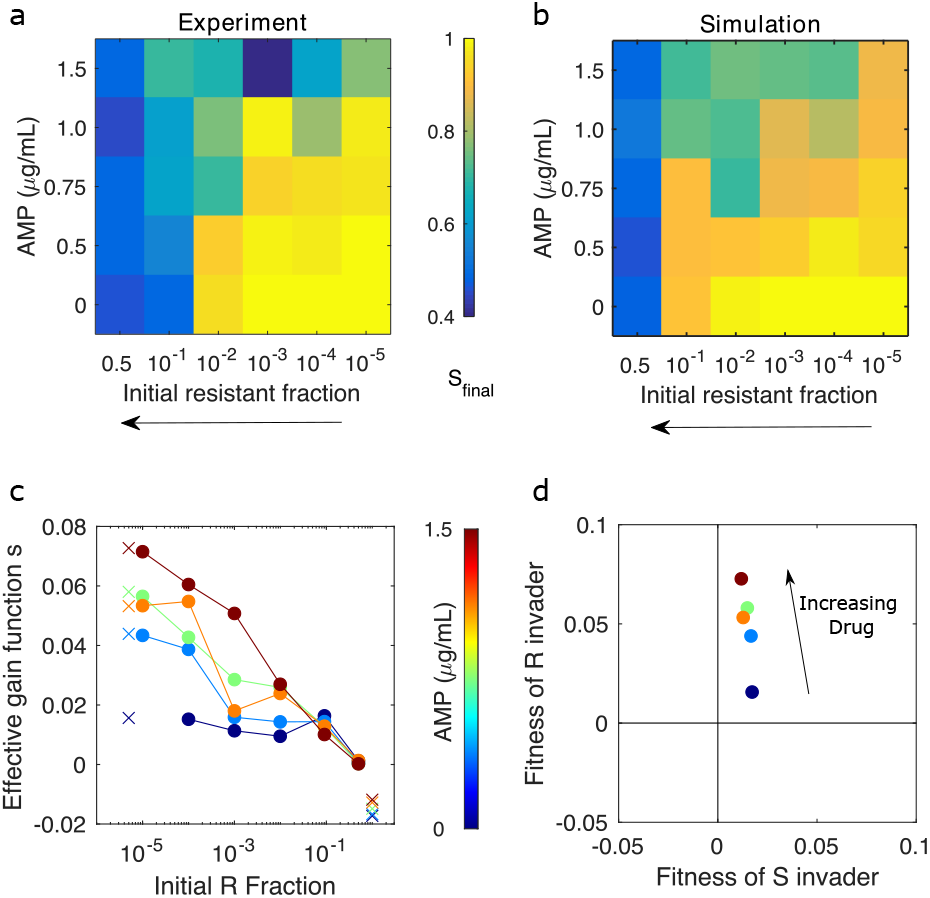
Experiments and simulations show that final composition depends on drug concentration and initial resistant fraction. Heat maps for experiment (a) and simulation (b) indicate final sensitive fraction (*S_final_*) for different values of AMP concentration and initial resistant fraction (which increases right to left, as in Figure 3). c) Initial and final fractions from experiment are converted to effective gain functions 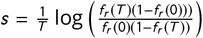, one for each drug concentration (blue, 0 to dark red, max). *f_r_(t)* is the fraction of resistant cells at time *t* (*t* = 0; *t* = *T* = 144 hrs). Markers (x) indicate linear extrapolation of each gain functions to an initial fraction of zero (left) or initial fraction of one (right). d. Extrapolated gain functions provide estimates of the (relative) fitness of a single resistant invader to an all-sensitive population (vertical axis) and a single sensitive invader to an all-resistant population (horizontal axis). These relative finesses can also be interpreted in the context of evolutionary game theory, where they suggestion long-term coexistence between sensitive and resistant cells (see Supplemental Text I)

### Frequency- and drug-dependent selection for resistance

To quantify the cooperation observed between sensitive and resistant cells, we calculated an effective gain function (98) for each drug concentration (Figure 4c; see also Supplemental Text I). The gain function *s* measures the selection coefficient for resistant cells as a function of initial population composition (resistant fraction). In exponentially growing populations, *s* reduces to the difference in per capita growth rates of sensitive and resistant subpopulations in a mixture with a given starting composition; more generally, it can be viewed as an effective measure of selection–specifically, it converts measured changes in population composition–in this case, at the beginning and end of the experiment–to an effective growth rate difference (in units of 1/time) that would give rise to the observed changes over the experimental time period *T*. We observe strong frequency- and drug-dependent selection for resistance (Figure 4c, Supplemental Text I), with selection highest when drug concentration is large and resistant fraction small (i.e. negative frequency-dependent selection (6)). Intuitively, resistance is strongly favored in scenarios where protective effects of the enzyme are minimized. By contrast, as the number of resistant cells (and the number of enzymes) increases, or the concentration of drug decreases, protective effects become increasingly significant, minimizing the selection for resistance as sensitive cells benefit from the presence of their resistant neighbors.

The limits of the gain function for small and large resistant fractions provide insight into potential invasion dynamics of sensitive and resistant cells. Recent work in cancer shows that these limits also correspond to entries of a payoff matrix describing effective evolutionary games (99, 98). In our experiments, extrapolation of the gain functions (Figure 4C) suggests that resistant invaders harbor a positive fitness advantage that increases with drug concentration; by contrast, sensitive invaders posses a small and largely drug-independent fitness advantage. In evolutionary game theory, these trends suggest a “leader” game, which would lead to stable coexistence between sensitive and resistant cells, similar to what was observed for non-small cell lung cancers cultured with cancer-associated fibroblasts (98). While we urge caution in strictly interpreting this result–in particular, the fitness of sensitive invaders relies on an extrapolated value very close to zero, which could correspond to marginal stability rather than a single coexistence fixed point–the analysis provides a convenient quantitative measure of cooperation.

### Resistant colonies offer protective effects over length scales several times the size of a single colony

Our results suggest that resistant cells offer cooperative resistance to sensitive cells over relatively long length scales. To directly probe the scale of this interaction, we inoculated one sensitive-only population and one resistant-only population on BHI agar plates supplemented with super-MIC concentrations of ampicillin. The colonies were inoculated at a fixed separation distance, which we varied from 1 to 3 cm, and we then tracked the area of the sensitive colony (in total pixel count) each day (Figure 5). At these drug concentrations, sensitive populations are unable to grow in isolation; any growth therefore reflects protective effects of the nearby resistant colonies. As a control, we also performed identical experiments in the absence of drug. We found that sensitive populations separated by up to 2 cm exhibited substantial growth at the end of the 6 day period, reaching nearly half the size of sensitive colonies grown without drug. By contrast, colonies separated by 3 cm–despite showing growth for the initial 3 days–eventually collapsed. These results indicate that on the timescale of our experiments, resistant colonies exhibit protective effects that extend up to two centimeters– almost three times the size of a single microbial colony.

**FIG 5.**
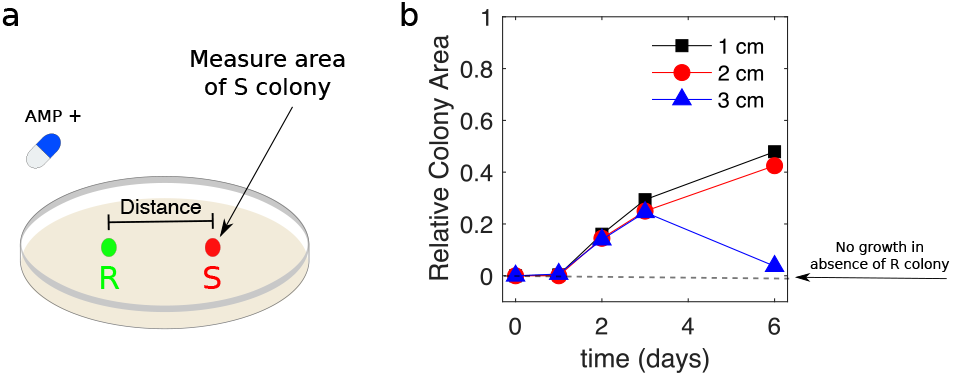
Sensitive colonies grow at super-MIC drug doses when separated from resistant colonies by multiple colony lengths. a. Resistant (green) and sensitive (red) populations are inoculated on day 0 at a fixed separation distance (1, 2, or 3 cm) on BHI-agar plates containing a super-MIC concentration of AMP (1.5 *μ*g/mL). The total area of the sensitive colony (in pixels) is measured daily. For comparison, an identical experiment is also performed in the absence of AMP. b. Area of the sensitive colony over time. Area is normalized by the area of sensitive colonies during a matched control experiment in the absence of drug. Colony area of one indicates growth is identical to that of sensitive colonies in the absence of drug. Note that in the absence of the resistant colony, the sensitive colony is not visible (dashed line).

### Inoculum density modulates segregation patterns and appearance time with little effect on expansion velocity

In addition to initial resistant fraction, the patterns of segregation are likely to be impacted by the initial density of the total population (i.e. the inoculum density) (100). To investigate this dependence, we grew colonies starting from initial densities that varied over 3 orders of magnitude both with and without super-MIC concentrations of drug. These densities correspond to initial populations containing, on average, between 20 (density of 10^−4^) and 2 × 10^5^ (density of 0.1) founder cells. Consistent with results in other (non-antibiotic) systems (100), we found that segregation is increased as initial density is decreased, both with and without drug, leading to patterns with thicker sectors (Figure 6). In addition, the expansion velocity was approximately constant across conditions, while arrival time decreased approximately logarithmically with initial density. If we assume that populations are growing at an effective exponential rate *k* and that arrival time corresponds to the population reaching some critical populations size (i.e. a visible threshold), the slope of the appearance time vs. log(density) plot provides an estimate of 1/*k*. For the conditions we measured, *k* ≈ 0.02 min^−1^, which corresponds to an approximate doubling time of 35 minutes, only slightly slower than drug-free growth in liquid media at the same temperature.

**FIG 6.**
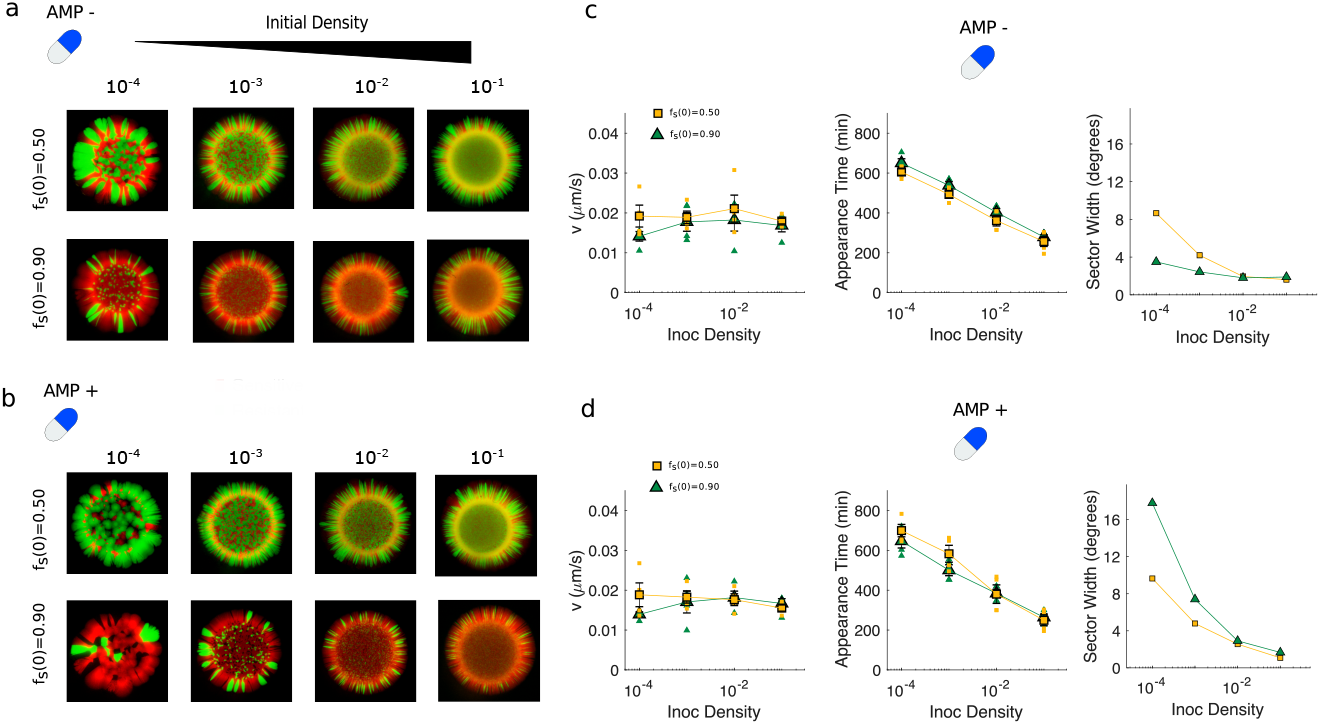
Decreasing inoculum density leads to increased segregation and faster appearance times but little change in expansion velocity. a. Spatial patterns for colonies starting from initial sensitive fractions of 0.5 (top) or 0.9 (bottom) and different starting (total) densities (increasing left to right) in the absence of AMP. b. Same as panel a, but in the presence of AMP at super-MIC concentrations (1 *μ*g/mL). c. Radial expansion velocity (left), appearance time (center), and median angular width of resistant sectors (right) for colonies grown under the same conditions as in panel a. Angular width is calculated using only the portion of the image outside of the homeland. Small points are individual replicates, large markers are mean over replicates ± standard error of the mean. Appearance time depends logarithmically on initial density, and the slope of the line is consistent with a per capita growth rate of approximately 0.02 min^−1^ (doubling time approximately 35 minutes). d. Same as panel c, but in the presence of super-MIC AMP.

### Initial populations unlikely to contain more than a single resistant cell can survive at super-MIC drug concentrations

We also investigated the combined effects of very small resistant fractions (< 10^−2^) and small inoculum densities, leading to extreme scenarios where the expected number of resistant cells in the initial population is much less than 1. Populations starting from 1 percent resistant cells expand at rates comparable to, but slightly less than, those observed for more resistant populations at the largest initial densities (> 10^−2^), though the arrival time of these colonies is significantly increased in the presence of drug (Figure 7a). For smaller inoculum densities, the colonies cannot be consistently tracked on the scanner, though microscopy images indicate coexistence of both sensitive and resistant cells. If we consider even smaller initial resistant fractions, we eventually reach a limit where growth is completely absent (Figure 7b)–results that are consistent with the fact that populations are unlikely to contain even a single resistant cells at inoculation. In this regime, the populations that do survive contain a single localized subpopulation of resistant cells, indicating that even a single resistant cell in the inoculum can provide sufficient protection to seed colony growth. Because these experiments are performed at super-MIC AMP concentrations, where sensitive cells are unable to survive in the absence of resistant cells, these local islands of resistance again suggest that protection is a long-range phenomenon that does not require sensitive and resistant cells to be in close spatial proximity.

**FIG 7.**
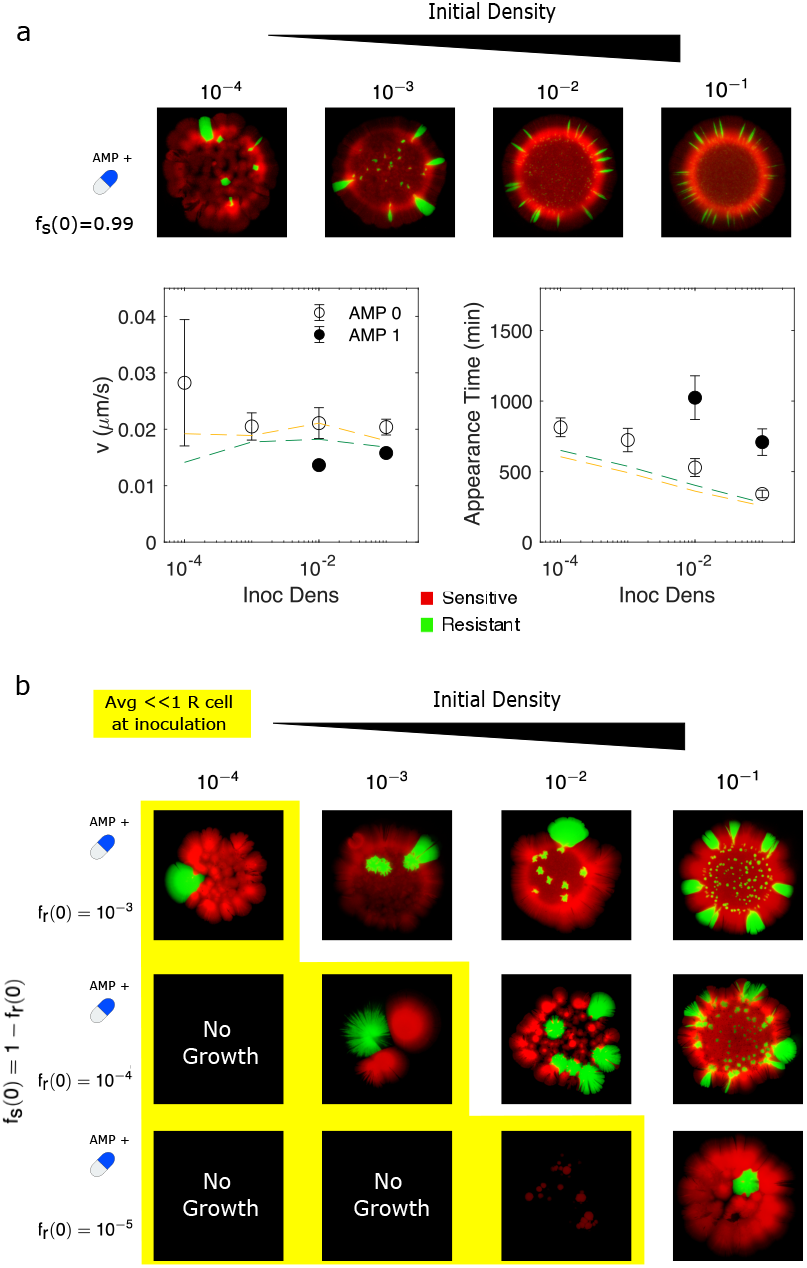
Initial populations unlikely to contain more than a single resistant cell are able to survive super-MIC concentrations. a. Colonies formed in the presence of super-MIC AMP (1 *μ*g/mL) starting from populations with one percent resistant cells at different total densities (left to right, corresponding to initial population sizes ranging from 10s to hundreds of thousands of cells). Bottom panels, expansion velocity (left) and appearance time (right) for colonies grown with (closed circles) and without (open circles) super-MIC AMP. Dashed lines (yellow, *f_s_*(0) = 0.5; green, *f_s_*(0) = 0.9) indicate mean trends from Figure 6 for comparison. Note that for initial densities less than 0.01, low colony density prohibits estimates of arrival time and expansion velocity on the scanner. b. Similar to panel a, but for increasingly smaller initial resistant fractions. Yellow region highlights conditions where initial population is unlikely to contain more than a single resistant cell (expected number 〈R_0_〉 ≪ 1). See also Figures S1-S2 for complete data set and drug-free controls.

### A simple biophysical model indicates that synergistic effects of multiple resistant cells contribute to long-range protection

Our results suggest that the protective effects of resistant cells can be relatively long-range, extending for many times the size of a single cell. Such long-range protection is expected in well-stirred communities without spatial constraints (5, 68, 11, 70) and is commonly observed on agar plates (78), especially when cells secrete the enzyme into the extracellular space. On the other hand, the protective effects of drug degrading enzymes appear considerably shorter-range in some naturally occurring isolates (18), where long-range protection may require alternative mechanisms such as persistence, and may depend on whether drug degradation is intracellular (66, 67, 14).

To investigate the variables that control this length scale, we developed a simple biophysical model of drug degradation that is assumed to occur at the surface of resistant cells (Supplemental Text II). In this model, which is based on classical models of diffusion-limited reactions (101) and cell receptor signaling (102), the resistant cell retains the enzyme, but protective effects can arise as freely diffusing drug molecules are degraded after coming into contact with the cell, which acts as a local reaction sink that reduces the local concentration of drug. For simplicity, we assume that diffusion of the drug occurs on a faster timescale than cell division, so the number and location of resistant cells is fixed. For a single resistant cell, the (steady state) radius *R_p_* of the protection zone–the region surrounding the resistant cell where drug concentration drops below some critical value *A_c_* (i.e. the MIC)–is given by

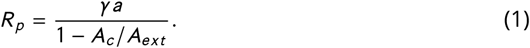

where *a* is the radius of the cell, *A_ext_* is the external drug concentration, and

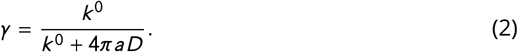

with *k*^0^ the effective rate constant for degradation when drug and enzyme are in close proximity, and *D* the diffusion constant of the drug in agar. When the rate of reaction is rapid (*k*^0^ ≫ 4*πaD*), the drug profile is determined by diffusion-limited reactions, with *γ* → 1 and the length scale of protection set by the radius a of the resistant cell. The diffusion-limited regime is an upper bound on the range of the protective effects offered by a single cell, and while it indicates that the protection zone increases rapidly as *A_ext_* → *A_c_* (near the MIC), the zone is limited by the size of the enzyme-producing cell. While β-lactamases are highly efficient enzymes (103) that frequently operate near the diffusion-limited regime (104, 105), protection zones would be further reduced by rate-reducing mutations in the enzyme (106, 105) or low expression levels (107, 108), both of which would effectively reduce *k*^0^. On the other hand, environmental factors that reduce diffusion of the drug–for example, the crowding that can occur in biofilms due to extracellular matrix (109)–drive the reaction toward the diffusion limited regime, even when local reaction rates are slower in dilute solutions.

These results suggest that a single resistant cell is unlikely to offer long-range protection across many cell lengths except when the drug concentration is very near the MIC. However, it is not clear whether the combined effects of multiple resistant cells might lead to significantly longer-range effects. To answer this question, we consider a spherical colony of radius *R* ≫ *a* that contains many cells (see Supplemental Text II). The colony includes *N_r_* enzyme-producing resistant cells uniformly distributed on its surface, and we assume that these resistant cells instantaneously degrade enzyme (one can easily relax this assumption to include finite reaction rates, as in Equation 1, but the goal here is to evaluate how effects of multiple cells combine, so we stick to the simplest case). For this idealized multi-cellular colony, the radius of protection is given by

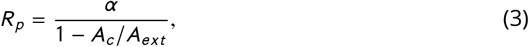

with

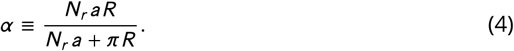

When the number of resistant cells is small (*N_r_* ≪ *R/a*), the length scale of the protection zone scales as *α* ~ *N_r_a*; that is, the net effect of *N_r_* resistant cells is approximately equal to *N_r_* times the effect of a single resistant cell. In the other extreme, as the number of resistant cells is large, *R_p_* approaches that for a purely resistant ensemble (*α* → *R*, see Supplemental Text II), and the range of protection is now set by the size of the entire cell ensemble. It is notable, however, that the effects accumulate nonlinearly as the number of resistant cells increases beyond a few, similar to the phenomenon first described for receptors on a cell surface (102). For example, in a spherical colony of *R* = 1000 microns containing resistant cells of radius *a* = 1 micron, the protection zone radius *R_p_* reaches 90 percent of its maximum value with only *N_r_* ≈ 3 × 10^4^. Remarkably, an *N_r_* of this size means that the colony surface contains relatively few resistant cells, covering only a small fraction 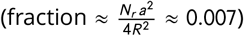 of the total surface area. As for cell receptors, the strong collective effects arise because drug diffusing near the colony surface is likely to make many contacts with the surface before diffusing away, dramatically increasing the probability of capture, even when the capture zones (resistant cells) are sparsely distributed on the surface.

This simple model indicates that the scale of the protective effects is set by the external drug concentration and the size of the colony. In a mixed population of sensitive and resistant cells, the protection zone grows rapidly with the number of resistant cells on the surface of the colony, producing a synergistic effect where the net effect of *N_r_* resistant cells is much greater than *N_r_* times the effect of a single cell. A similar argument applies to single cells with a finite number of enzymes bound to the cell surface. In each case, increasing the number of enzymes–whether by increasing expression levels in a single cell (e.g. plasmid vs chromosome-based resistance), or by increasing the number of resistant cells in a community–is expected to significantly increase the range of protective effects, even when the surface area of the colony is dominated by sensitive cells and enzyme-free regions. The key, however, is that the resistant cells should be uniformly distributed across the surface. In reality, clonal growth dynamics and selection are coupled, creating clustered regions of resistant cells, at least under some conditions, that would reduce this synergy. The sensitivity of the protection zone to these geometric and biophysical variables may partially explain the wide range of resistance length scales observed in different microbial systems (66, 67, 68, 11, 69, 14).

## DISCUSSION

Our results complement a number of recent studies showing that antibiotic resistance in microbial communities can reflect cooperative interactions between resistant and sensitive cells (5, 7, 10, 11, 12, 20, 13, 14, 15, 16, 17, 70). The genetic segregation characteristic of many spatially-extended microbial systems often leads to scenarios where “producer cells", those cells that supply a public good such as a drug-degrading enzyme, preferentially benefit other producers, maintaining cooperation and discouraging exploitation by non-producers (39, 23). While we observe the formation of spatial sectors of sensitive or resistant cells, particularly in larger populations, we also observe widespread survival of sensitive cells in mixed populations, even at drug concentrations lethal to sensitive-only populations. The ubiquity of these non-producers can be partially explained by the presence of long-range cooperative interactions, where resistant cells offer a protective effect that extends far beyond their neighboring cells– extending for up to several centimeters on the timescales of our experiments. It is perhaps surprising to see such long-range effects, particularly because *β*-lactamase producing *E. faecalis* are generally not motile and are not believed to release the enzyme into the extracellular space (86, 88). Cooperation is therefore unlikely to result from long-range diffusion of the enzyme itself, but instead from diffusion of drug molecules to spatially fixed “sinks” (resistant cells) where they are degraded. In systems where the cells are motile or the enzyme can freely diffuse, the dynamics of cooperation may be considerably more complex and even, in some limits, mimic behavior of classic reaction-diffusion (or even sub-diffusive) systems (110, 111). For example, recently-developed algorithms (112) draw on analogies with electrodynamics to calculate profiles of chemical species (in this case, antibiotic) in the presence of multiple sinks (resistant cells). These algorithms could be applied to the irregular geometries observed in colony growth experiments–relaxing the assumption that resistant cells are uniformly distributed–to calculate protection zones in more realistic scenarios. These dynamics could prove even more rich in synthetic or multi-species communities, where metabolic co-dependencies (19) or rational design strategies (113) may lead to additional intercellular interactions.

It is important to keep in mind several limitations of our study. From a technical perspective, our study relies on fluorescent imaging of multi-species colonies. We used the observed segregation patterns to estimate population composition, but one should interpret these estimates with some caution, as they do not capture composition as accurately as more labor-intensive methods such as flow cytometry, plate-based CFU estimates, or confocal microscopy at single-cell resolution. While we find that these estimates are not particularly sensitive to the thresholds used in our image analysis (see Figure S3), we note that the analysis is unlikely to capture fine-scale changes in composition, particularly in well mixed regions or for colonies with very narrow segments. Therefore, in all cases, we also provide the colony images themselves, and the composition estimates should be viewed as convenient, if imperfect, metrics for quantifying the major trends as a particular parameter (e.g. drug concentration) is varied.

Perhaps most importantly, this work is entirely *in vitro* and therefore does not capture the complex environmental factors that may drive the formation of biofilms– and the expansion of resistance–in clinical scenarios. In particular, the range and magnitude of cooperative interactions may depend on specific conditions–dictated, for example, by diffusion constants and growth rates characteristic of a given host environment–potentially leading to different dynamics (5, 68, 11, 70, 66, 67, 14). In fact, our models suggest that the length scales of cooperation may vary significantly with even relatively small changes in biophysical parameters, such as enzyme expression level, because the net effect of multiple reaction sinks (enzymes or resistant cells) can be strongly synergistic, and that synergism itself depends on the degree of uniformity in the spatial distribution of cells. In addition, our experiments take place on relatively short timescales, where population behavior is likely dominated by selection of existing resistant strains rather than evolution of new resistance mutations. Our results may not, therefore, reflect dynamics that are stable on evolutionary timescales, where stochastic appearance of new phenotypes may lead to entirely new behavior. Finally, we note that our primary goal in developing simple mathematical models was to probe the effects of local vs global cooperative resistance in a spatially extended community, and to gauge the effects of different biophysical parameters on those effects. The models neglect many known features of the antibiotic response in living microbial communities, including pharmacological details of drug activity, phenotypic heterogeneity, and persistence. The biophysical models also depend on idealized systems with simple geometries, while the patterns observed experimentally are considerably more variable. Our hope is that future work will build on these results using increasingly detailed models, similar to those developed for other systems (112, 114, 115), and experimental tools for directly measuring spatial distribution of drug (116, 117), which may help to describe even the fine-scale structure of the observed spatial patterns. Despite the simplicity of our models, they reproduce observed population-level qualitative trends and, along with experiments, support the idea that interactions between resistant and sensitive *E. faecalis* communities may promote survival of drug sensitive cells on length scales the size of entire communities.

## ACKNOWLEDGMENTS

This work is supported by the National Science Foundation (NSF No. 1553028 to KW), the National Institutes of Health (NIH No. 1R35GM124875-01 to KW), and the Hartwell Foundation for Biomedical Research (to KW). We would like to thank Dr. Wen Yu for her initial work in troubleshooting the document scanner system for tracking colony growth, and members of the Wood lab (UM) for helpful discussions. The format for this preprint is adapted from the American Society for Microbiology (ASM) template available on Overleaf.com.

## CONFLICT OF INTEREST

The authors declare that they have no conflict of interest.

## Notes

### Competing Interest Statement

The authors have declared no competing interest.

### Summary of Updates

This version includes a simple biophysical model of diffusion-limited drug degradation, additional quantitative analyses of the observed selection dynamics, and minor corrections for style.

