## Supplemental Text for "Spatial segregation and cooperation in radially expanding microbial colonies under antibiotic stress"

### SUPPLEMENTAL TEXT I: COOPERATIVE RESISTANCE AS AN EVOLUTIONARY GAME.

Recent work has shown that collective dynamics in cancerous tumors can be described using experimentally-parameterized models of effective evolutionary games (98). The approach is powerful, in part, because it can be applied using relatively simple experimental measurements that track population composition. Here we apply the framework from (98) to the cooperative resistance observed in our experiments. The analysis indicates that our system sits near the boundary of two qualitatively distinct dynamical regimes, though extrapolation of our data suggests a leader game resulting in stable coexistence between sensitive and resistant cells (though we cannot rule out marginal stability that would occur at the boundary). The analysis provides a convenient quantitative measure of cooperation (via entries of an effective payoff matrix) with a simple and intuitive interpretation. While a full elucidation of the game dynamics will require further study (most notably, time-dependent measures of composition across conditions), we anticipate this preliminary analysis will serve as a foundation for future investigations of cooperative resistance under different conditions.

The idea, briefly, is to assume that populations of each cell type grow with a per capita rate ($g_i$) of the form,

$$\begin{aligned} g_R &= Af_r + B(1-f_r) \\ g_S &= Cf_r + D(1-f_r) \end{aligned} \tag{S1}$$

where $g_R$ is the growth rate for resistant cells, $g_S$ the rate for sensitive cells, $A, B, C, D$ are coefficients of an effective payoff matrix, and $f_r$ is the fraction of resistant cells. Competition assays between sensitive and resistant cells are used to infer the parameters of the payoff matrix, which fully determine the phase space of possible dynamical regimes: stability of purely resistant populations, stability of sensitive populations, bistability (where both types of homogeneous populations are possible, depending on initial conditions), and coexistence (98). More specifically, the model reduces to a replicator-like equation for $f_r$

$$\frac{df_r}{dt} = f_r(1-f_r)G(f_r) \tag{S2}$$

where $G(f_r)$ is a gain function that depends on differences in the payoff matrix entries

$$G(f_r) = g_R - g_S = (B-D)(1-f_r) - (C-A)f_r. \tag{S3}$$

The steady state dynamics depend only on the sign of the differences $B-D$ and $C-A$, which correspond to the relative fitness of a resistant invader and the relative fitness of a sensitive invader, respectively.

Kaznatcheev *et al* show that the payoff matrix can be estimated by quantifying the frequency dependence of sub-population growth rates at different (initial) population compositions (98), where the parameters correspond to the $f_r \to 0$ and $f_r \to 1$ intercepts of the single species data (as in Equation S1). While we do not have detailed time series data on growth rates for individual sensitive and resistant populations, we can apply a slightly coarser approach by estimating the gain function $G(f_r)$ using initial and final measurements of population composition ($f_r$) estimated from fluorescence data. Specifically, we calculate an effective gain function $s$–similar to the selection coefficients often used to quantify pairwise competition experiments–as

$$s = \frac{1}{T}\log\left(\frac{f_r(T)(1-f_r(0)))}{f_r(0)(1-f_r(T))}\right) \tag{S4}$$

where $T$ is the total length of the experiment (approximately 6 days). Equation S4 is just a two-point estimate of $G(f_r)$ which assumes that the dynamics between time 0 and time $T$ are determined by Equation S2 with $G(f_r) = s$ a constant (which, however, will vary for each experiment that starts with a different initial value of $f_r$). By estimating $s$ for each experiment (each starting from a different composition), we can estimate the effective gain function, and the limits of that function (extrapolated to $f_R = 0$ and $f_R = 1$) provide estimates of $C - A$ and $B - D$, respectively.

### SUPPLEMENTAL TEXT II: BIOPHYSICAL MODEL OF PROTECTIVE RESISTANCE

To estimate the length scale for protective resistance, we first consider a single, fixed, spherical, enzyme-producing (resistant) cell of radius $a$ placed at the origin. Our goal is to find the spatial profile of antibiotic concentration $A(r)$ a distance $r$ from the origin, assuming that $A(\infty) = A_{ext}$ is the external concentration far away from the resistant cell and the drug follows the diffusion equation,

$$\frac{\partial A}{\partial t} = D\nabla^2 A, \tag{S5}$$

where $D$ is the diffusion constant of drug. The surface of the cell contains $N$ bound copies of the enzyme, each of which degrades nearby antibiotic according to Michaelis Menten kinetics at rate

$$\text{rate} = \frac{k^0_{kat} A(a)}{A(a) + K^0_m} \approx k^0 A(a) \tag{S6}$$

where $k^0 \approx k^0_{cat}/K^0_m$ and the superscripts indicate that these are local rates valid when antibiotic is present at the cell surface $r = a$ (i.e. these rates do not include the effects of diffusion of antibiotic to the enzyme). The last step in Equation S6 assumes $A(a) \ll K^0_m$, though it is straightforward to incorporate the full nonlinear rate equation as needed.

**Infinitely rapid degradation at the cell surface.** When the number of enzymes ($N$) is large–so that the entire surface of the cell is covered with enzyme–and the reaction rate ($k^0$) is infinitely fast, the concentration of drug is vanishingly small at the cell surface. This scenario corresponds to the classic Smoluchowski problem (101) from chemical kinetics. The steady state solution of Equation S5 with $A(\infty) = A_{ext}$ and $A(a) = 0$ is given by

$$A(r) = A_{ext}\left(1 - \frac{a}{r}\right). \tag{S7}$$

For a given profile $A(r)$, the size $R_p$ of the "protection zone"–the region surrounding the resistant cell for which $A$ is less than some critical value $A_c$ (i.e. the MIC)–is given by

$$R_p = \frac{a}{1 - A_c/A_{ext}}. \tag{S8}$$

For very large drug concentrations $A_{ext} \gg A_c$, the protection zone $R_p$ includes only the resistant cell ($R_p \to a$). However, as $A_{ext}$ is reduced to $A_c$, the protection zone grows rapidly, potentially encompassing many cell lengths. For example, if $A_{ext} = 1.1A_c$ (i.e. the concentration exceeds the MIC by only 10 percent), the protection zone $R_p = 11a$.

**Finite degradation rates at the cell surface.** Equation S8 is an upper bound on the size of the expected protection zone for a single cell; it is valid when the enzyme reaction rate at the cell boundary is sufficiently fast that only diffusion of the drug to the cell limits degradation. While β-lactamases are known to be highly efficient enzymes that often approach diffusion-limited rates in solution, there are likely examples where degradation is slower because of limitations to the intrinsic turnover rate or, perhaps, low expression of the enzyme, leading to a non-zero concentration $A(a)$ of drug near the cell surface. In this case, the antibiotic profile reduces to

$$A(r) = A_{ext}\left(1 - \gamma\frac{a}{r}\right), \tag{S9}$$

with $\gamma = \frac{A_{ext}-A(a)}{A_{ext}}$ a factor containing the dependence on $A(a)$, which itself can be related to the local reaction rate constant $k^0$ by assuming a partially reflecting boundary condition at the cell surface (101)

$$k^0 A(a) = 4\pi a^2 D \frac{\partial A}{\partial r}\Big|_{r=a}, \tag{S10}$$

leading to

$$A(a) = \frac{4\pi a A_{ext} D}{k^0 + 4\pi a D}. \tag{S11}$$

and

$$\gamma = \frac{k^0}{k^0 + 4\pi a D}. \tag{S12}$$

In the limit $k^0 \gg 4\pi a D$, we have $\gamma \to 1$ and the profile reduces to the diffusion-limited case (Equation S7). On the other hand, when the reaction rates are finite and of the same order as the diffusion effects, the protection zone $R_p$ will be reduced by a factor $R_p \to \gamma R_p$. Unfortunately, estimates of $k^0$ (the reaction rate given that enzyme and substrate are localized together) are not typically available. The fact that many β-lactamases show near diffusion-limited rates means that $k^0 \gg 4\pi r_E D$, with $r_E$ the reaction radius of a single enzyme and $D$ the diffusion constant of drug in solution. However, the radius of a single cell is at least 2-3 orders of magnitude larger ($a \gg r_E$), so the diffusion-limited regime for a single enzyme is not necessarily diffusion-limited when the enzymes sit on the surface of a cell. More specifically, covering a cell of radius $a$ with a diffusion-limited enzyme (i.e. sufficiently large $k^0$ to yield $A(r_E) = 0$ at the reaction radius $r_E$ for a single enzyme) does not necessarily mean that the concentration $A(a)$ at the surface of that cell will be zero. On the other hand, diffusion of drug in a bacterial colony may be significantly slower than diffusion in solution, which would favor reduced concentrations at the cell surface.

**Synergistic protection in cell ensembles with multiple resistant cells.** The upper bound for the length scale of the protection zone surrounding a single resistant cell is set by $a$, the radius of the cell. For drug concentrations on the order of 2x the MIC–similar to those used in our experiments–one would only expect protection to extend for a few cell lengths, but experiments suggest protection extends considerably farther in cellular communities. One explanation could be that the collective effects of multiple resistant cells are synergistic, producing a protection zone that reflects more than a simple accumulation of single cell effects.

To investigate this possibility, we consider a large spherical ensemble of radius $R \gg a$ that contains many resistant cells. For simplicity, we assume that each resistant cell expresses a sufficient quantity of enzyme to completely degrade any drug that reaches the cell surface. In this case, the concentration of drug approaches zero at the surface of the ensemble, and the problem is again equivalent to the Smoluchowski problem, but the length scale is now set by $R$

$$A(r) = A_{ext}\left(1 - \frac{R}{r}\right). \tag{S13}$$

A large ensemble of resistant cells is therefore expected to exhibit protection zones with length scales set by the size of the ensemble. As before, this serves as an upper bound, and finite reaction rates / rapid diffusion may decrease this range.

Surprisingly, however, these large-scale protection zones can be achieved even when the ensemble contains only a small fraction of resistant cells. Specifically, let us assume the ensemble is made up of both sensitive and resistant cells. Furthermore, assume a total of $N_r$ resistant cells, each represented by a circle of radius $a$, are uniformly distributed over the surface of the sphere. The total fraction of the ensemble surface covered by resistant cells is given by $f \approx \frac{N_r a^2}{4R^2}$. How does the size of the protection zone depend on this fraction (or equivalently, on $N_r$)?

To solve the problem, we exploit the analogy with electrodynamics used in (102) in the context of diffusion-limited cellular signaling. Specifically, the steady state diffusion equation $D\nabla^2 A = 0$ has the same form as the equation for the electrostatic potential $\phi$ in a charge-free space, $\nabla^2\phi = 0$. As a result, the diffusive current density is analogous to the electric field vector, and the total diffusive current $J$ entering a closed surface is analogous to the total charge $Q$ on a surface. The condition $A = 0$ at the spherical surface is equivalent, in the electrodynamics analog, to the surface being isopotential. The analogy allows us to equate the total diffusive flux $J$ through any closed surface to the electrical capacitance $C$ (in cgs units) of an isolated conductor of the same shape (102) according to

$$J = 4\pi C D A_{ext}. \tag{S14}$$

This analogy is particularly convenient since the capacitance, $C$, has been calculated for conductors of many different sizes and shapes. In the case of our spherical ensemble of cells, the capacitance of interest is that of an insulating sphere of radius $R$ covered uniformly with $N_r$ conducting disks of radius $a$, all connected with infinitesimal wires to form a single conductor. The capacitance $C$ of this object is derived in (102) as

$$C = \frac{N_r a R}{N_r a + \pi R}, \tag{S15}$$

an expression that is valid when the distance between neighboring disks (in this case, resistant cells) is large compared to the radius of a single disk (cell). The total diffusive flux for the cell ensemble is therefore given by

$$J = J_{max} \frac{N_r a}{N_r a + \pi R}, \tag{S16}$$

where $J_{max} = 4\pi R D A_{ext}$ is the flux through a spherical ensemble containing only resistant cells. Assuming the flux is uniform over the sphere (no angular dependence), the drug profile outside of the ensemble ($r \geq R$) is

$$A(r) = A_{ext}\left(1 - \frac{\alpha}{r}\right) \tag{S17}$$

with

$$\alpha \equiv \frac{N_r a R}{N_r a + \pi R} \tag{S18}$$

Similarly, the size of the protection zone becomes

$$R_p = \frac{\alpha}{1 - A_c/A_{ext}}. \tag{S19}$$

When the number of resistant cells is small ($N_r \ll R/a$), the length scale of the protection zone scales as $\alpha \sim N_r a$; that is, the net effect of $N_r$ resistant cells is approximately equal to $N_r$ times the effect of each cell. In the other extreme, as the number of resistant cells is large, $R_p$ approaches that for a purely resistant ensemble ($\alpha \to R$). But interestingly, the effects accumulate rapidly (and nonlinearly) as the number of resistant cells increases beyond a few (as in the case for cell receptors on the surface of a single cell (102)). As an example, for a spherical ensemble of $R = 1000$ microns, the protection zone radius $R_p$ reaches 90 percent of its maximum value with only $N_r \approx 30000$, corresponding to $f \approx \frac{N_r a^2}{4R^2} \approx 0.007$–that is, when less than 1 percent of the surface is covered with resistant cells.

### SUPPLEMENTAL FIGURES

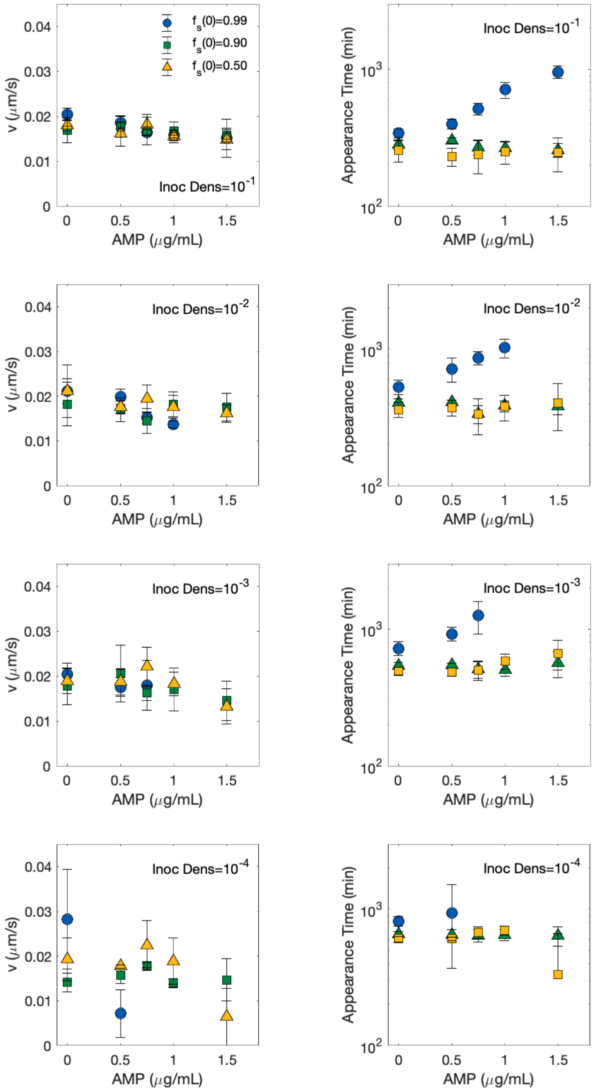


**FIG S1** Expansion velocity (left) and appearance time (right) of mixed colonies at different concentrations of ampicillin (AMP). Each row corresponds to a particular inoculum density. We note that particularly at low inoculum density, it becomes difficult to track colony expansion because the colonies do not have well defined boundaries on the scanner images. At these densities, we cannot rule out weak concentration dependence of expansion velocity.

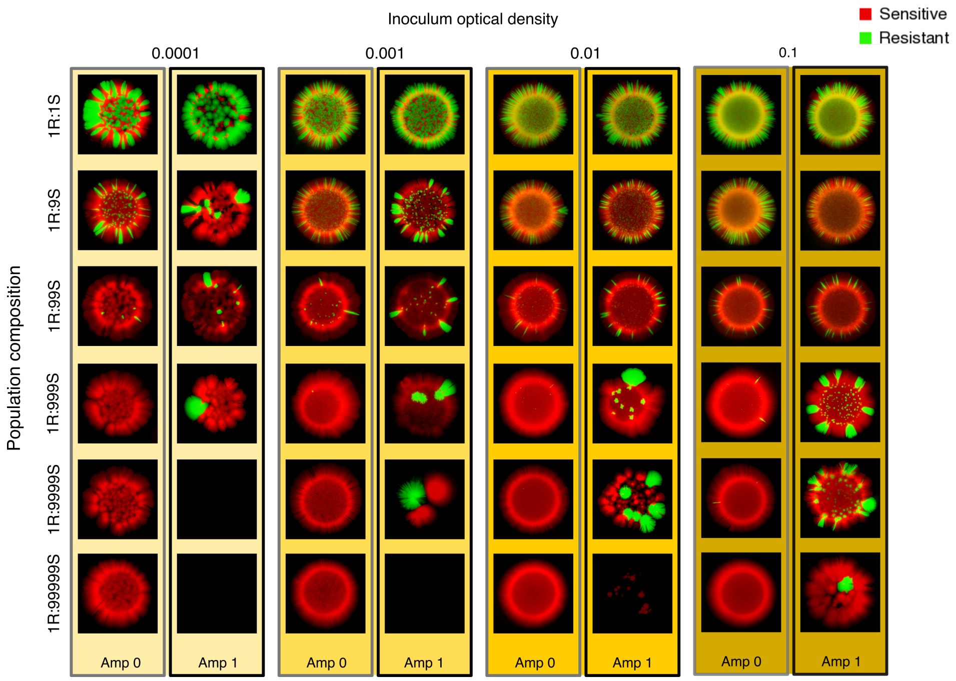


**FIG S2** Spatial patterns for example colonies starting from different initial densities (columns) and different initial compositions (rows). For each condition, colonies were grown in the presence or absence of super-MIC AMP (1 *mu*g/mL).

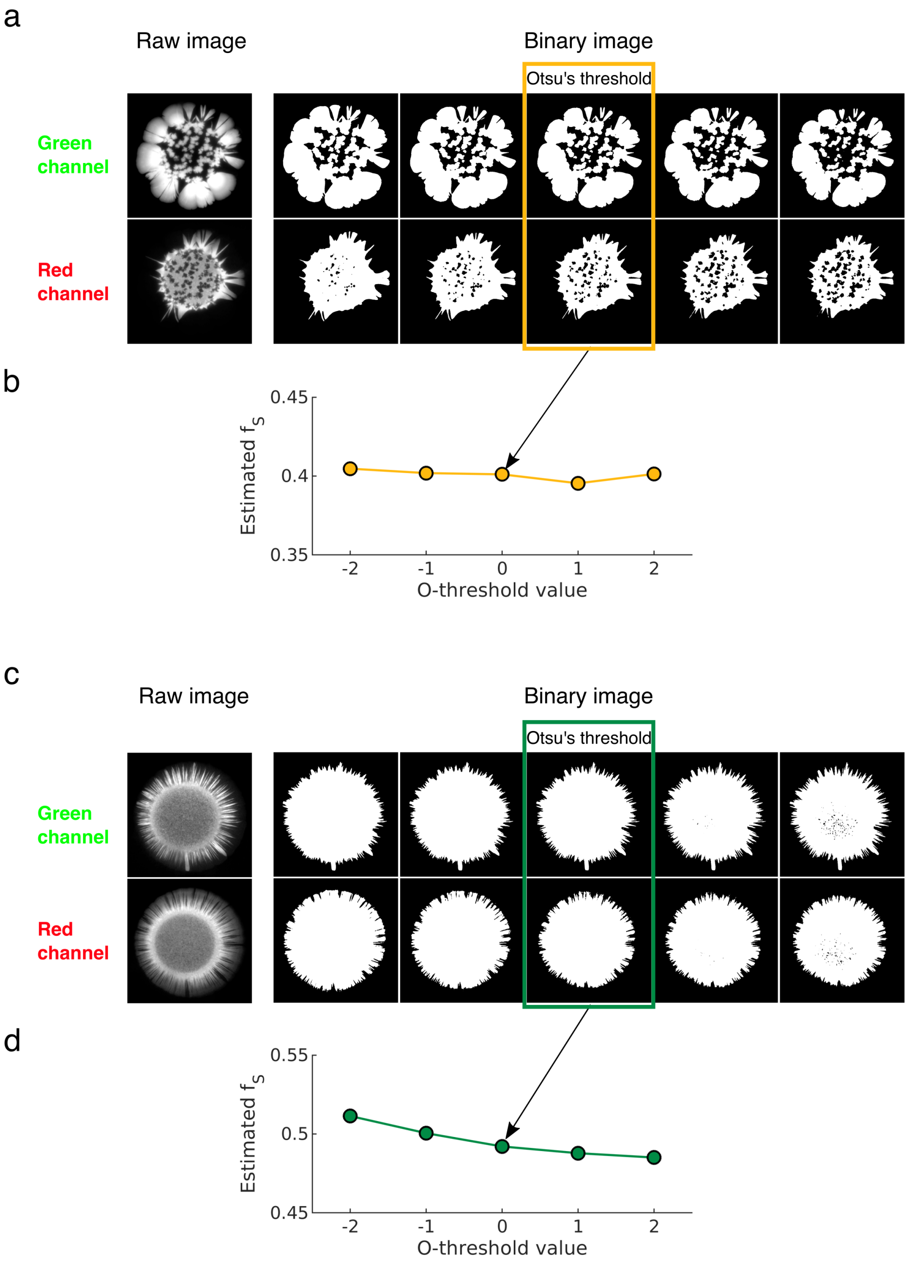


**FIG S3 The estimated fraction is insensitive to the thresholding process** a. Raw images of green and red channel acquired by imaging the mixed colonies (left) and binary image for different threshold (right). Middle images show the binarization using Otsu's method and other images show ± 25% ,± 12.5% change in the threshold. b. The fraction of sensitive cells estimated from images with different threshold. c and d. Similar to panel a and b, but for colonies with narrow monoallelic regions, which makes the binarization difficult and thereby prone to over/underestimation of cells.
